# Unconventional structure and function of PHD domains from Additional Sex Combs-like proteins

**DOI:** 10.1101/2024.12.08.627434

**Authors:** CJ Reddington, AR Walsh, C Göbl, PD Mace

**Affiliations:** Biochemistry Department, School of Biomedical Sciences, University of Otago, Dunedin, New Zealand; Mātai Hāora — Centre for Redox Biology and Medicine, Department of Pathology and Biomedical Science, University of Otago, Christchurch, New Zealand; School of Biological Sciences, University of Canterbury, Christchurch, New Zealand

## Abstract

The Polycomb Repressive-Deubiquitinase (PR-DUB) complex removes ubiquitin from Lysine residue 119 on histone H2A (H2AK119Ub) in humans. The PR-DUB is composed of two central protein factors, the catalytic Breast Cancer type 1 susceptibility protein (BRCA1)-activating protein 1 (BAP1), and one of Additional Sex Combs-like 1–3 (ASXL1–3). A Plant Homeodomain (PHD) at the C-terminus of ASXL proteins is recurrently truncated in cancer, and was previously proposed to recognise epigenetic modifications on the N-terminal tail of histone H3. Here we demonstrate that the ASXL PHD domain lacks features required for histone tail recognition and is unable to bind histone H3 epigenetic marks. Modelling the structure of the ASXL PHD using AlphaFold3 suggests that the domain has an atypical fold and that the isolated ASXL PHD can chelate a single Zinc ion *in vitro*, compared to the two ions conventionally bound by PHD domains. Recently, the ASXL PHD was shown to bind an auxiliary set of PR-DUB interactors, named methyl CpG-binding domain proteins 5 (MBD5) and 6 (MBD6). We show that the ASXL PHD-MBD5 and -MBD6 complexes are stable *in vitro,* and surprisingly contain a composite Zinc-binding site at the interface between the two proteins. Overall, this data suggests an unconventional pairing of domains coordinate key functions of the PR-DUB — a non-canonical PHD domain from ASXL proteins obligately partners with MBD5 and 6, which were themselves misannotated because they cannot bind to methylated DNA.

## Introduction

Chromatin, the structure used to efficiently store DNA, is formed with DNA wrapped twice around an octamer of histone proteins to form a nucleosome. Two copies of four histone proteins, H2A, H2B, H3 and H4, interact within the nucleosome core. The quaternary structure of the nucleosome forces the N- and C-terminal tails from histone proteins to protrude for covalent chemical modification (Chen & Dent, 2014). Attachment of ubiquitin to histone H2A at K119 limits gene expression (Cao & Yan, 2012). Ubiquitin is attached to H2AK119 by the Really Interesting New Gene (RING) E3 ubiquitin ligase Polycomb Repressive Complex 1 (PRC1, Cohen et al., 2020) and is removed by the PR-DUB (Reddington et al., 2020; Scheuermann et al., 2010).

The PR-DUB has regulatory functions in the cell cycle, cellular development and DNA damage response, and determines short-term changes to gene expression (reviewed in Di Croce & Helin, 2013; Mozgova & Hennig, 2015; Parreno & Martinez, 2022; Schuettengruber et al., 2017). The core of the human PR-DUB is composed of a catalytic deubiquitinase, BAP1, and one of three regulatory ASXL proteins (mutually exclusive, interchangeable). The BAP1 deubiquitinase is comprised of two conserved, functional domains; the Ubiquitin C-terminal Hydrolase (UCH) confers histone protein deubiquitination, whereas the UCH L5-like domain (ULD) interacts with the Deubiquitinase-associated Domain (DEUBAD) of ASXL proteins (Daou et al., 2015; Sahtoe et al., 2016; Scheuermann et al., 2010). Interaction with the ASXL DEUBAD is required to form a composite ubiquitin-binding site, shown by crystallization of the *Drosophila* PR-DUB in 2018, and later in 2019 (De et al., 2019; Foglizzo et al., 2018). Recently, the structure of the human PR-DUB bound to ubiquitinated nucleosome was resolved by cryo-electron microscopy. In complex with chromatin, BAP1 and ASXL are positioned to restrict the PR-DUB toward removal of H2AK119Ub and not other histone marks (Ge et al., 2023; Thomas et al., 2023). ASXL1–3 also contain a DNA-binding, HB1, ASXL, Restriction Endonuclease-Helix-Turn-Helix (HARE-HTH) domain at their N-terminus and a C-terminal PHD finger, thought to bind histone modifications (Figure 1a, Aravind & Iyer, 2012).

**Figure 1.**
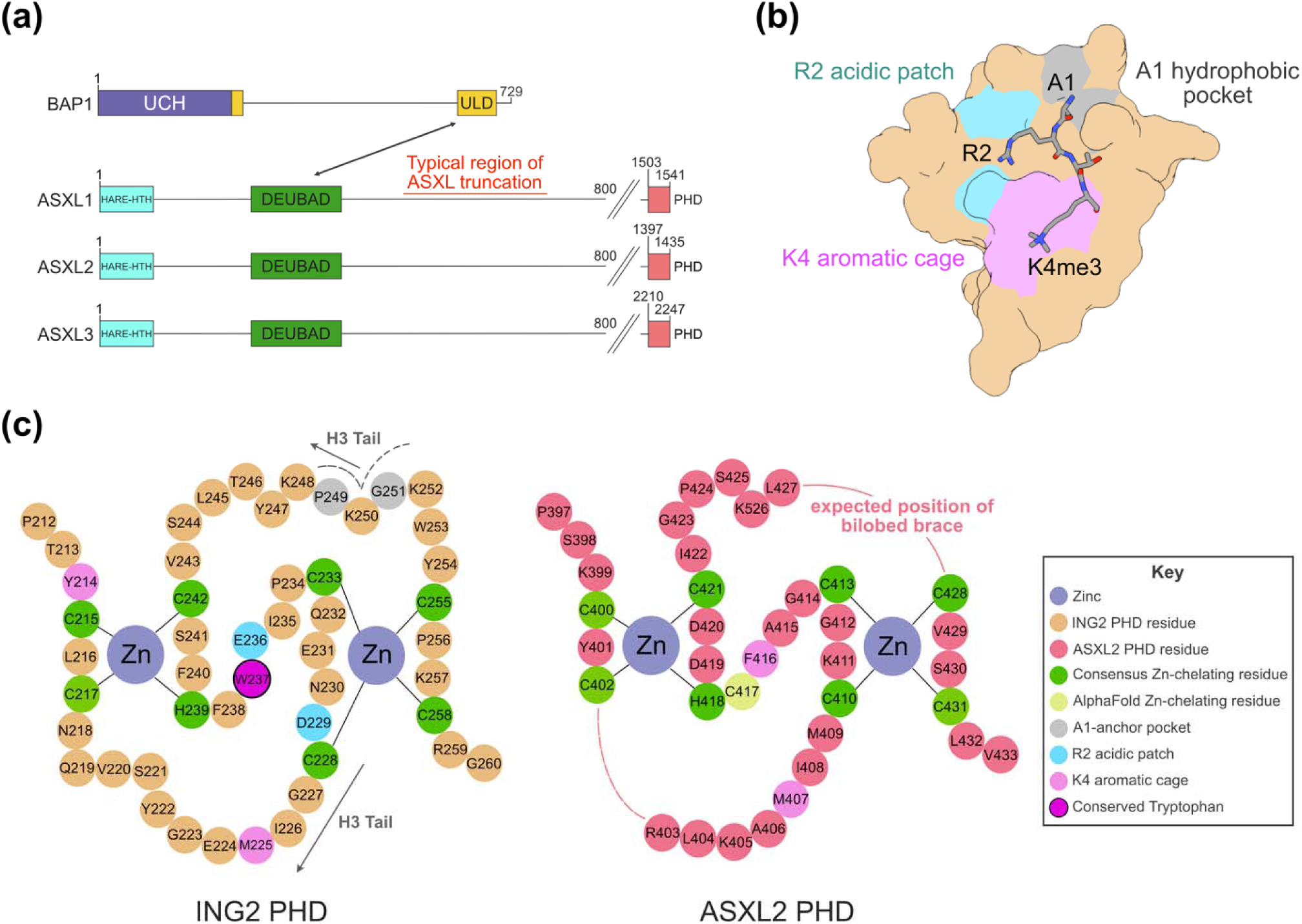
Atypical organisation of the ASXL PHD domain. **(a)** Schematic representing domain structure of human BAP1 and ASXL1–3. Functional regions are labelled, with the ASXL DEUBAD and BAP1 ULD domains shown to interact (arrow). The region typically afflicted by truncating ASXL mutations is highlighted (red). Residue boundaries are given for the PHD domains of ASXL1–3. **(b)** Structure of histone H3K4me3 peptide (residues 1–4) bound to the PHD domain of ING2 (residues 212–263, PDB: 2G6Q). The ING2 PHD structure is shown in surface representation and colour-coded for key features as in Figure 1c. The histone H3K4me3 peptide is shown in stick representation and coloured by element. **(c)** Organisation of residues from the PHD domains of ING2 (left, residues 212– 260) and ASXL2 (right, residues 1397–1433, listed as residues 397–433). Functional residues are highlighted according to the key (inset). The position of the histone H3 tail is shown in dark grey, shortened regions on the ASXL PHD are shown as lines.

ASXL1–3 are recurrently truncated in human hematological malignancies, particularly Acute Myeloid Leukaemia. ASXL mutations typically occur in the last exon and truncate the protein, causing loss of the PHD domain (Figure 1a, Balasubramani et al., 2015). Often, ASXL truncation products are expressed, escape nonsense-mediated decay (Inoue et al., 2016) and interact to form aberrant versions of the PR-DUB (Asada et al., 2018; Balasubramani et al., 2015; Daou et al., 2018). The mutant PR-DUB has increased stability and deubiquitinase activity, facilitating widespread removal of H2AK119Ub (Asada et al., 2018; Daou et al., 2018). Given the ASXL PHD is repetitively lost in cancer, the domain may have an important functional role. This is further reiterated by the observation that similar inherited truncations of ASXL1 (Bohring Opitz Syndrome, Hoischen et al., 2011), ASXL2 (Shashi-Pena Syndrome, Jiao et al., 2022) and ASXL3 (Bainbridge-Ropers Syndrome, Yang et al., 2020) cause developmental disorders.

PHD domains conventionally possess a ‘binuclear treble-clef’ Zinc finger fold, where Cysteine and Histidine residues chelate two structurally integral Zinc ions in a Cys^4^-His-Cys^3^ structural motif (Dodd et al., 2004; Kaur & Subramanian, 2016; Li & Li, 2012). Typically, PHD domains bind the terminal Alanine (A1), Arginine (R2) and Lysine (K4) residues of the histone H3 N-terminal tail to selectively recognise histone modification state (Figure 1b). Most commonly, PHD domains bind either unmodified, di- or tri-methylated H3K4, where H3K4me2 and H3K4me3 specify active gene expression states (Figure 1b, Li & Li, 2012; Sanchez & Zhou, 2011; Slama & Geman, 2011). The histone methyl specificity of the ASXL PHD is not currently clear (Aravind & Iyer, 2012; Park et al., 2016). Here, we use a variety of methods to show the ASXL PHD domains are unlikely to bind modified or unmodified histone tails. Instead, AlphaFold modelling and subsequent validation with recombinant protein showed that the domain has an atypical fold. Through study of the recently-characterised interaction between the ASXL PHD domains and Methyl CpG-binding domains 5 (MBD5) and 6 (MBD6, Tsuboyama et al., 2022) we found the MBD proteins form a stable complex with the ASXL PHD, and that an additional Zinc-binding site is shared between them. Accordingly, MBD5 and MBD6 appear to be essential to PR-DUB function and their loss as interaction partners is likely causative in ASXL truncation-driven disease.

## Methods

### Plasmids and cloning

Expression vectors were generated by Ligation Independent Cloning (LIC, Luna Vargas et al., 2011). All insert DNA was synthesised by Integrated DNA Technologies (IDT). Typically, insert fragments were flanked by 5’-CAGGGACCCGGT-3’ (forward) and 3’-TAACCGGGCTTCTCCTCG-5’ (reverse) overhangs for LIC cloning. For bacterial protein expression, DNA fragments were cloned into expression vectors from the NKI LIC suite (Luna Vargas et al., 2011). DNA sequences for the Inhibitor of Growth 2 (ING2, residues 199–261), ASXL2 (residues 1375–1435) and *Drosophila* Additional Sex Combs (Asx, residues 1610–1668) PHD domains all were cloned into an N-terminal Glutathione S-transferase (GST)-3C protease fusion vector (addgene 108710) for expression. The DNA sequence for the ASXL2 PHD domain (residues 1395-1435) was also cloned into an N-terminal His_6_-SUMO3-SENP fusion vector (addgene no.) for expression alone, and with the MBD domain of MBD6 (residues 17-81) for co-expression, connected by a DuetLinker sequence. For expression alone, the MBD6 MBD domain (residues 17-81) was cloned into an N-terminal StrepII-3C fusion vector (addgene 108707).

### Bacterial protein expression and purification

Proteins were expressed in *Escherichia coli (E. coli)* BL21 (DE3) cells using Luria Broth (Formedium), with cells grown at 37 °C in a shaking incubator. Once cultures met the optical density desired (0.5–0.8 OD_600_), they were transferred to 18 °C for overnight (O/N) growth. Protein expression was induced by addition of isopropyl β-D-1-thiogalactopyranoside (IPTG) to a final concentration of 0.2 mM. For proteins that bind Zinc, ZnCl_2_ was added to a final concentration of 0.1 mM. Bacterial cell pellets were resuspended in a buffer of 300 mM NaCl, 20 mM Tris pH 8.0, before being frozen at −20 degrees. Cell pellets were thawed for lysis and diluted in a buffer of 300 mM NaCl, 50 mM Tris pH 8.0, 10% (*v/v*) glycerol, 10% (*v/v*) sucrose, 10 mM imidazole. Hen-egg lysozyme (Hampton Research) was added to a final concentration of 166 µg/mL and incubated on ice for 30 minutes. Later, cells were disrupted either by repeated rounds of sonication (Sonifier Cell Disruptor, Bronson Sonic Power Co.) or emulsification (Emulsiflex, Avestin).

All GST-fusion proteins were purified with soluble protein fractions bound to Protino Glutathione Agarose resin 4B (GSH resin, Macherey-Nagel). For Isothermal Titration Calorimetry (ITC), the PHD domains of ING2, ASXL2 and Asx were bound to GSH resin in batch before washing three times in 300 mM NaCl, 50 mM Tris pH 8.0, 10% (*v/v*) glycerol, 10% (*v/v*) sucrose, 10 mM imidazole. Bound protein was cleaved using GST-fused 3C protease O/N, with Dithiothreitol (DTT) added to a final concentration of 2 mM. Cleaved PHD domain proteins were flowed through a Poly-Prep Chromatography hand-column (Bio-Rad) before being purified by size-exclusion (SE) chromatography on an ÄKTA Pure 150 Chromatography system, using a Superdex 75 Increase 10/300 column (GE Healthcare). A buffer of 300 mM NaCl, 20 mM HEPES and 0.5 mM Tris (2-carboxyethyl) phosphine (TCEP) was used for SE chromatography purification of ITC proteins, with eluent fractions studied by Sodium Dodecyl Sulfate-Polyacrylamide Gel Electrophoresis (SDS-PAGE, Sup. Fig. 1). Fractions containing proteins of interest were concentrated by centrifugal ultrafiltration before being snap-frozen and stored at −80 °C.

Proteins for histone H3 enzyme-linked immunosorbent assay (ELISA) and the initial Zinc-binding analysis were expressed as GST-fusions, purified and eluted from GSH resin through a Poly-Prep Chromatography hand-column (Bio-Rad) by addition of fresh glutathione-containing buffer (300 mM NaCl, 50 mM Tris pH 8.0, 10 mM glutathione). Proteins were purified by SE chromatography on an ÄKTA Pure 150 Chromatography system, using a Superdex 75 Increase 10/300 column (GE Healthcare) in a buffer of 300 mM NaCl, 20 mM HEPES (2 mM DTT for ELISA samples, no reducing agent for Zinc assay preparations). Protein fractions were studied by SDS-PAGE, with fractions containing proteins of interest combined, concentrated by centrifugal ultrafiltration, snap-frozen and stored at −80 °C (Sup. Fig. 2, 5).

For the second Zinc-binding analysis, SUMO2-fusion proteins with Hexa-histidine (His_6_) tags were purified from HIS-Select Nickel (Ni^2+^) Affinity Gel resin (Sigma) through Poly-Prep Chromatography hand-columns (Bio-Rad). The soluble protein fractions were added to equilibrated Ni^2+^ resin before washing three times in 300 mM NaCl, 50 mM Tris pH 8.0, 10% (*v/v*) glycerol, 10% (*v/v*) sucrose, 10 mM imidazole. Proteins were eluted from the column by the addition of 300 mM imidazole-containing buffer (300 mM NaCl, 50 mM Tris pH 8.0, 10% (*v/v*) glycerol, 10% (*v/v*) sucrose, 300 mM imidazole). The StrepII-fusion MBD6 protein was purified and eluted from Strep-Tactin®XT 4Flow® high capacity resin (IBA Lifesciences) through a Poly-Prep Chromatography hand-column (Bio-Rad). Following the addition of the soluble protein fraction, the resin was washed three times in 1x Buffer W (100 mM Tris/HCl, pH 8.0, 150 mM NaCl, 1 mM EDTA) and eluted by the addition of 1x Buffer BXT to the column (100 mM Tris/HCl, pH 8.0, 150 mM NaCl, 1 mM EDTA, 50 mM biotin) (IBA Lifesciences). Proteins were purified by SE chromatography on an ÄKTA Pure 150 Chromatography system, using a Superdex 75 Increase 10/300 column (GE Healthcare) in a buffer of 300 mM NaCl, 20 mM HEPES. Protein fractions were studied by SDS-PAGE, with fractions containing proteins of interest snap-frozen and stored at −80 °C (Sup. Fig. 7).

### Isothermal titration calorimetry

ITC was performed in a low volume TA Affinity ITC instrument (TA instruments). Histone peptides — constituting unmodified histone H3 (residues 1–8, AS-65152), H3K4me2 (residues 1–10, AS-64370-1) and H3K4me3 (residues 1–10, AS-64371-1) — were sourced from AnaSpec and diluted to 400 μM in the buffer used to purify all input proteins (300 mM NaCl, 20 mM HEPES, 0.5 mM TCEP). Histone peptides were placed in the ITC sample syringe prior to analysis. PHD domain proteins were assayed at 60 μM in the ITC sample cell, with the input concentration determined by Bicinchoninic acid (BCA) assay. The performance of the ITC instrument was measured in runs of water against water, with a heat of injection of less than 2.5 ± 2 μJ allowed. For ITC, twenty 2.5 μL peptide injections were performed, with the sample cell set to 25 °C, 75 rpm spinning rate. To process binding energies, baselines were calculated using NITPIC and thermodynamic parameters by global analysis in SEDPHAT (Brautigam et al., 2016). All data was plotted in GUSSI (Brautigam et al., 2016). Blank runs (Histone peptides into buffer) were collected but not subtracted through experimental heat shifts (no titration of binding observed, Sup. Fig. 1).

### Histone H3 modification ELISA

The Pre-Sure Histone H3 Peptide Array ELISA Kit (Colorimetric) from EpiGenTek (P-3104-96) was used to probe binding of a broad set of Histone H3 epigenetic marks by the PHD domains of ASXL2, Asx and ING2. The organisation of modified Histone peptides on the Histone H3 ELISA is detailed in Sup.Table 1. Proteins were assayed at 200 ng/well, diluted in the wash buffer provided (pH 7.2 to 7.5). Protein concentration was determined by BCA assay, with 50 μL added per well. Plates were sealed in parafilm, with solutions incubated for 2 hours at 4 °C (gently rocked). All subsequent wash steps were performed using the working wash buffer provided, with three initial 5-minute washes. Next, the ELISA was incubated with 50 μL α-GST primary antibody (Rabbit monoclonal, 1:500 dilution, Invitrogen #700775) for one hour at 37 °C. Three further 5-minute washes were followed by addition of a horse-radish peroxidase (HRP)-fused α-Rabbit IgG secondary antibody (Goat polyclonal, 1:5000 dilution, Abcam ab6721-1). After washing, bound protein signal was collected by addition of 100 uL detecting solution. The detecting solution was incubated for 7 minutes before 100 uL stopping solution was added to wells. Plate absorbance was measured at 450 nm on a Multiskan GO plate scanner (Thermo Scientific). The blank sample reading was subtracted through all samples, with the mean and standard deviation of the binding signal calculated in Microsoft Excel, and data graphed in GraphPad Prism.

### Structural modelling in AlphaFold3

All structural models were predicted on the AlphaFold3 server (Abramson et al., 2024; Jumper et al., 2021). Residue boundaries and the number of Zinc ions that were modelled to bind each protein are described where relevant in the main text.

### Nuclear Magnetic Resonance (NMR)

The sample was prepared using uniform ^15^N labelling through expression in minimal medium containing ^15^NH_4_Cl as the sole nitrogen source. After purification, the protein solution was concentrated to 42.5 μM and transferred into a 50 mM Phosphate buffer, pH 6.5. 50 μl D_2_O was added as a lock reagent yielding an overall volume of 500 μl prior to measurement of ^1^H^15^N HSQC experiments. Spectra were measured at 298 K on a Bruker 600 MHz spectrometer with an Advance III console and a TXi triple-resonance probe including z-axis gradients and a trim pulse during the first INEPT transfer for solvent suppression. 128 complex data points were measured in the indirect dimension by accumulating 80 scans with an inter-scan delay of 1 second. The data were acquired using Topspin 3.6.5, processed using NMRPipe (Delaglio et al., 1995) and analyzed using CCPNMR 2.4.2 (Vranken et al., 2005).

### Zinc-binding assay

The protocol for Zinc detection followed the method outlined in Doyle *et al*., 2019. The input concentration of proteins was determined by Bradford assay, where protein samples were not diluted prior to analysis. All proteins were purified in a buffer of 300 mM NaCl, 20 mM HEPES without reducing agent, with a target input concentration of 20 μM. To denature proteins, an 8 M stock of guanidine hydrochloride (GdnHCl) was prepared, with 337.5 μL GdnHCl added to 50 μL quantified input protein. The GdnHCl-protein mixture was incubated for 40 minutes at room temperature, inverted five times every 10 minutes. For detection of released Zinc, a 4X 4-(2-pyridilazo)resorcinol (PAR, Sigma-Aldrich) working solution (400 μM) was made from a 10 mM PAR stock (diluted in 222 mM HEPES, pH 8.0). 112.5 μL of 4X PAR was added to the Gdn-HCl-protein mixture before sample absorbance was measured at 500 nm. The concentration of released Zinc was calculated from a PAR-Zn^2+^ absorption coefficient of 66000 M^-1^cm^-1^ and adjusted for dilution. The performance of the Zinc-binding assay was tested by comparison to serial dilutions of a 200 μM ZnCl_2_ stock.

## Results

### The sequence of the ASXL PHD argues against canonical PHD domain function

Typically, PHD domains bind terminal residues of the histone H3 N-terminal tail (A1, R2 and K4) through conserved elements of the PHD domain fold (Figure 1B, Li & Li, 2012; Sanchez & Zhou, 2011; Slama & Geman, 2011). The residues that specify histone-tail recognition are well understood and can be identified on the primary sequence of a classical PHD (Slama & Geman, 2011). The organisation of residues within a conventional PHD domain are displayed for Inhibitor of Growth 2 (ING2) on the left-hand side of Figure 1c. For comparison, the amino acid sequence of the ASXL2 PHD (residues 1397–1433) is modelled alongside (Figure 1c, right). While the Cys^4^-His-Cys^3^ Zinc chelation motif typical to PHD domains is evident for the ASXL PHD (Aravind & Iyer, 2012; Li & Li, 2012), most other residues that function in histone binding are missing. Noticeably, the ‘bilobed brace’, specific to the N-terminal Alanine (A1) of histone H3 (solid red line, Aravind & Iyer, 2012), and a conserved Tryptophan, characteristic to PHD domains that bind H3K4 methyl marks (purple, outlined, Bortoluzzi et al., 2017), are missing (Aravind & Iyer, 2012). The PHD domains of ASXL1–3 also do not contain an obvious aromatic cage — required to bind histone methyl modifications — with few aromatic amino acids conserved (pink, Bortoluzzi et al., 2017; Li & Li, 2012; Slama & Geman, 2011).

Therefore, prior reports of histone methyl-binding (Aravind & Iyer, 2012; Park et al., 2016) are contradictory to the sequence features readily identifiable in the ASXL PHD domains. To fully explore whether the ASXL PHD can bind histone marks, we experimentally probed histone modification recognition by the ASXL PHD domain.

### The ASXL PHD is not able to recognise common histone H3 epigenetic modifications

The amino acid sequence of the PHD domains between ASXL1–3 is highly conserved (89% identity between ASXL1 and ASXL2, 86% identity for ASXL1 and ASXL3), but the ASXL2 PHD domain gave substantially greater yields when expressed as a GST-fusion protein in *E. coli*, so was used for *in vitro* binding studies. In addition, the PHD domain of Asx — the sole relative of ASXL proteins from *Drosophila melanogaster* (residues 1610–1668, Foglizzo et al., 2018; Scheuermann et al., 2010) — was also tested. To initially determine whether the ASXL PHD would bind histone H3K4 methyl marks, isothermal titration calorimetry (ITC) was performed. Peptides comprising unmodified histone H3, H3K4me2 and H3K4me3 (Figure 2a) were titrated into samples of the ASXL2, Asx or ING2 PHD domains, with the ING2 PHD acting as a positive control (Freyer & Lewis, 2008; Pierce et al., 1999). Previously, Tryptophan fluorescence spectroscopy was used to measure affinity of the ING2 PHD to H3K4me2 (1.5 +/- 1 µM) and H3K4me3 (15 +/- 4 µM) *in vitro* (Peña et al., 2006). Binding affinity of the ING2 PHD measured using ITC was comparable to reported values from fluorescence measurements, with dissociation constants of 4.2 and 8.2 µM for H3K4me2 (Sup. Fig. 1), and 4.1 and 6.5 µM for H3K4me3 (Figure 2a, Peña et al., 2006). In contrast, experimental analysis of H3K4me2, H3K4me3 and unmodified histone H3 peptide-binding showed no evidence of binding for the ASXL2 or Asx PHD, suggesting these protein domains are incapable of binding to unmodified histone H3, or H3K4 methyl marks under these conditions (Figure 2a, Sup. Fig. 1)

**Figure 2.**
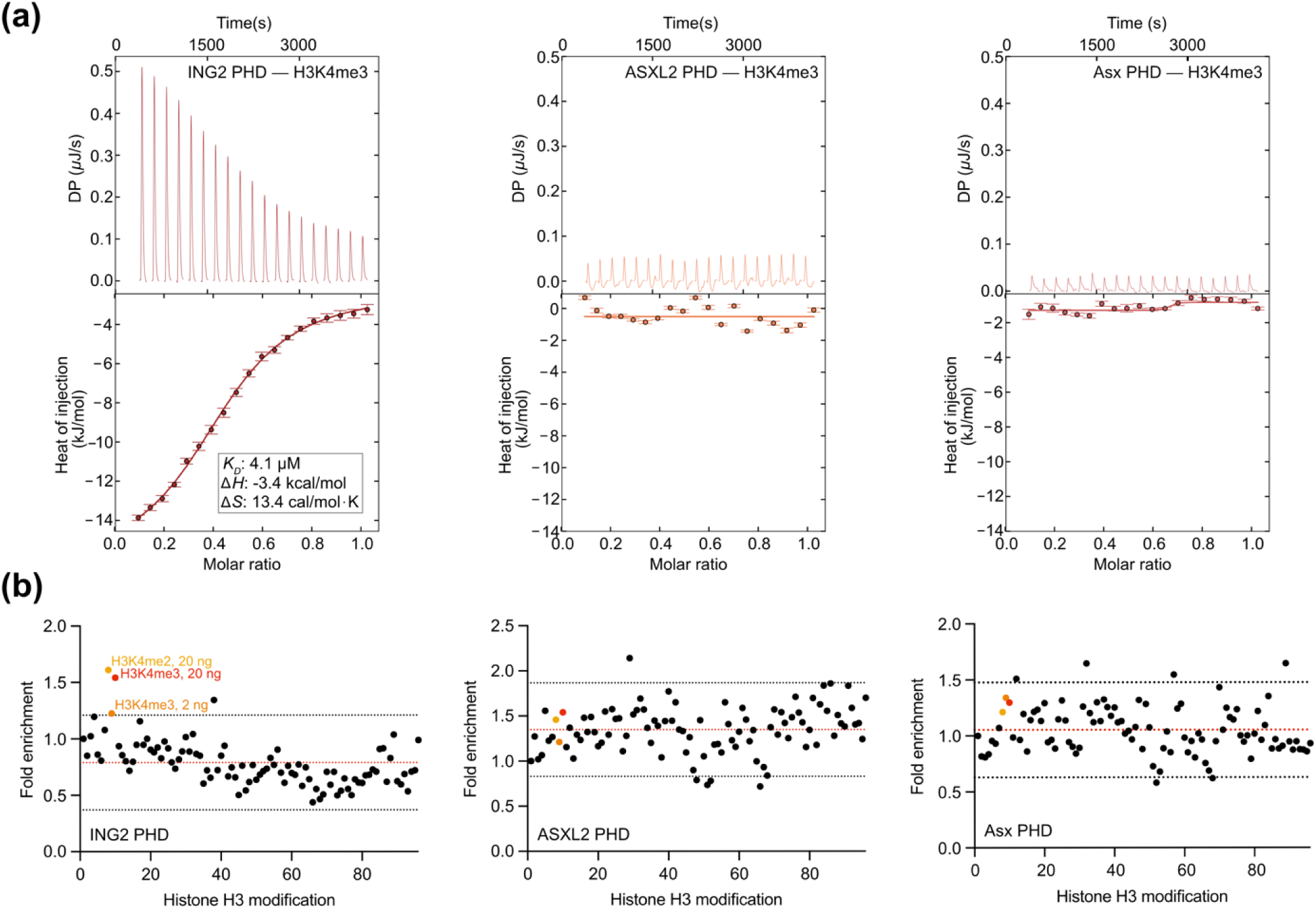
The ASXL PHD domain is unable to bind common histone H3 modifications. **(a)** ITC for interaction of the ING2 (residues 199–261), ASXL2 (residues 1375–1435), and Drosophila Asx PHD (residues 1610–1668) domains with peptides of H3K4me3 (residues 1–10). Key binding data are listed for the ING2 PHD titration. **(b)** Fold enrichment of protein signal collected in representative histone H3 ELISA experiments, titrated with samples of the ING2 (residues 199–261), ASXL2 (residues 1375–1435), and Drosophila Asx PHD (residues 1610–1668) domains. Single points correspond to each of the 96 peptides spotted on the ELISA plate, in the order displayed in Appendix Table 1. Values are derived from the fold-enrichment of HRP signal relative to blank. The mean (red) and two standard deviations from the mean (black) are dashed lines. Peptides of H3K4me2 (20 ng) and H3K4me3 (2, 20 ng) are highlighted yellow, orange, and red, respectively (labelled).

To probe whether the ASXL PHD may bind other histone H3 epigenetic modifications, a commercially manufactured enzyme-linked immunosorbent assay (ELISA) of modified histone H3 peptides was tested. Duplicate ELISA experiments were performed with the PHD domains of human ING2 and ASXL2 (Figure 2b, Sup. Fig 2). A single replicate was collected for the *Drosophila* Asx PHD (Figure 2b). As anticipated, the ING2 PHD showed preference for binding to methylated histone H3K4, where H3K4me2 and H3K4me3 were enriched by more than twice the standard deviation from the mean in replicate experiments (yellow, orange, red in Figure 2b, Peña et al., 2006). While there was some variability in this ELISA-based measurement, only H3K4me2 and H3K4me3 were bound in duplicate and in a concentration dependent manner (Figure 2b, Sup. Table 1). In contrast, ELISAs for the PHD domains of ASXL2 and Asx showed no consistent enrichment, where no histone mark was enriched in a concentration dependent manner (Figure 2b, Sup. Table 1). Our *in silico* characterization of PHD-determining residues (Figure 1c), ITC analysis of histone H3K4 methyl marks (Figure 2a) and histone H3 ELISA (Figure 2b) data all argue against histone modification-binding for the ASXL PHD. Thus, we propose the ASXL PHD is unable to bind histone H3 epigenetic marks and does not function conventionally as a PHD domain.

### The ASXL PHD is divergent from classical binuclear treble-clef domains

An assortment of protein domains bind Zinc to increase stability (Gamsjaeger et al., 2007; Krishna et al., 2003). Of these, several chelate two Zinc ions through the ‘binuclear treble-clef’ fold predicted for the ASXL PHD. Common binuclear treble-clef domains include the PHD, RING, ‘LIN-1, Isl-1 and MEC-3 (LIM) and Myeloid Translocation Protein 8, Nervy and DEAF1 (MYND) Zinc fingers, which each possess a distinct structure, function and fold (Dodd et al., 2004; Kaur & Subramanian, 2016; Matthews et al., 2009). Typically, the primary sequence of related domains are compared to determine their relationship (Kaur & Subramanian, 2016). However, the ASXL PHD appears unusual, does not function as a traditional PHD domain and may have been misannotated by this approach.

In the current work, several attempts were made to derive a macro-molecular structure for the ASXL PHD. The domain, however, proved too small and flexible to produce crystals for X-ray crystallography and weakly-dimerised with itself to limit study by NMR (data not shown). Instead, we predicted the structure of the ASXL2 PHD (residues 1399–1435) bound to two Zinc ions using AlphaFold3 (Figure 3a, Sup. Fig. 3, Abramson et al., 2024; Jumper et al., 2021). Overall, predicted Local Distance Difference Test (pLDDT) scores for the PHD domain were moderate (Sup. Fig. 3). AlphaFold assembled the ASXL2 PHD as a short, globular domain with an antiparallel β-sheet adjacent to an α-helix (Figure 3a). Unexpectedly, the Zinc-binding site predicted at the N-terminus of ASXL2 PHD (labelled, Figure 3a) did not involve Cys1400 (bold, Figure 3a). Cys1400 has conventionally been predicted to bind Zinc within the ASXL PHD (Figure 1c, Aravind & Iyer, 2012), forming part of the Cys^4^-His-Cys^3^ structural motif that is typical to PHD domains (Bienz, 2006; Li & Li, 2012). The adjacent Cysteine and Histidine residues at positions 1417 and 1418 were otherwise modelled to bind Zinc at the ASXL2 PHD N-terminus in an overall square-planar orientation (yellow, Figure 1c, 3b). Zinc ions typically are coordinated by proteins in tetrahedral fashion (Kuppuraj et al., 2009), which is clearly evident within the ASXL2 PHD C-terminal Zinc-binding site in Figure 3c.

**Figure 3.**
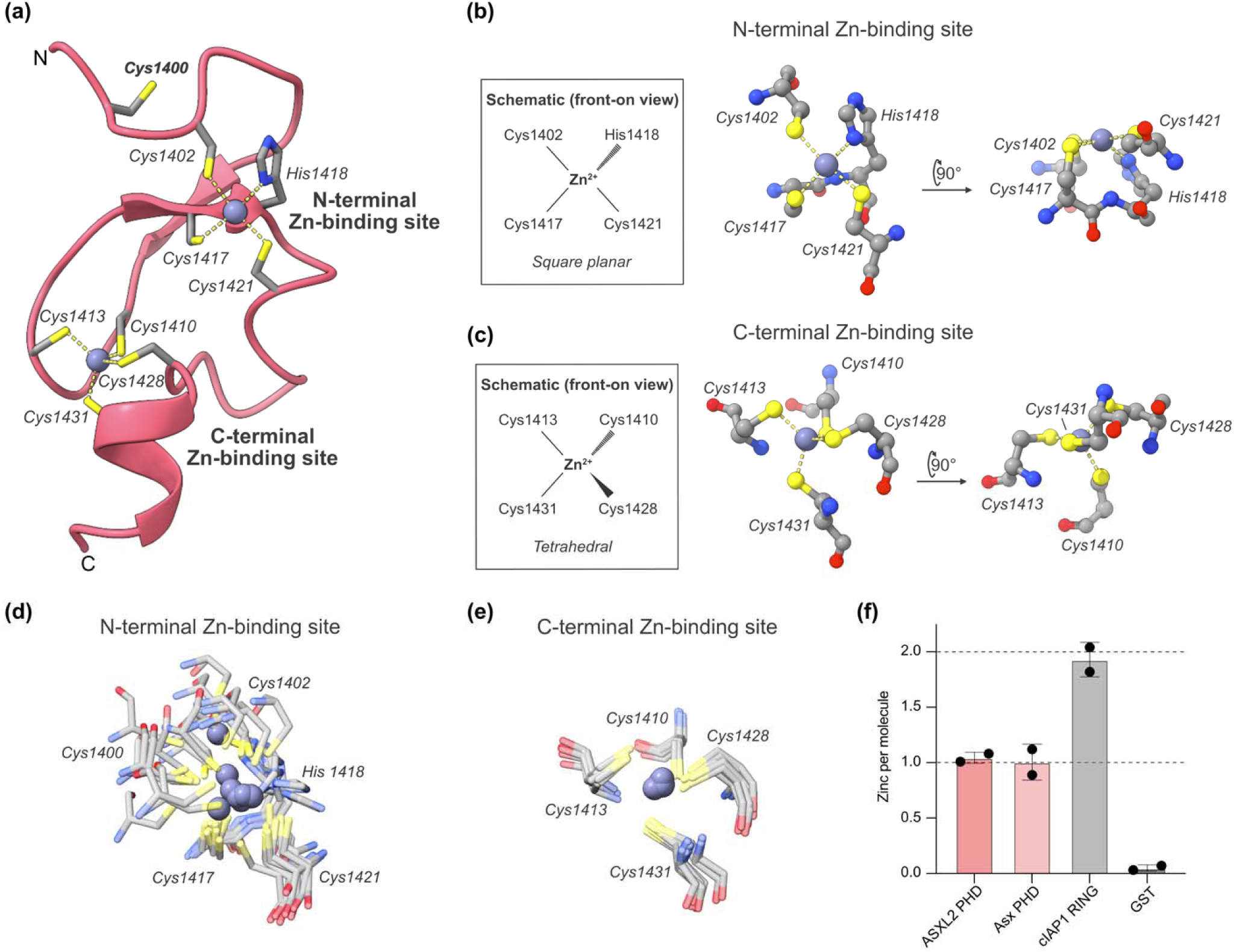
The N-terminal Zinc-binding site of the ASXL PHD is poorly formed and unable to chelate Zinc. **(a)** Structure of the ASXL2 PHD domain (residues 1399–1435) predicted using AlphaFold3, and modeled to bind two Zinc ions. The predicted structure of the ASXL2 PHD is shown in cartoon representation (pink). Residues predicted to bind Zinc are labeled, shown in stick representation and colored by element (Cys1400 labeled in bold). The N- and C-terminal Zinc-binding regions are labeled, respectively. **(b)** The square planar, N-terminal Zinc-binding site modeled for the ASXL2 PHD, shown from the perspective in Figure 3a. A schematic (left) illustrates the position of chelating residues in the central image (ball and stick representation). The perspective is rotated 90 degrees to give the view on the right. **(c)** The tetrahedral, C-terminal Zinc-binding site modeled for the ASXL2 PHD, shown as in Figure 3c. **(d)** The N-terminal Zinc-binding site of the ASXL2 PHD, modeled ten times in AlphaFold3 (residues 1399–1435). Relevant Zinc-chelating residues are shown as translucent sticks. The variably modeled position of Zinc within the domain is highlighted (silhouette, opaque). **(e)** The C-terminal Zinc-binding site of the ASXL2 PHD, consistently modeled in AlphaFold3 (ten models, residues 1399–1435), shown as in Figure 3d. **(f)** Zinc-binding assay for the ASXL2 (residues 1375–1435) and Asx (residues 1610–1668) PHD domains, compared to the RING domain of cIAP1 (residues 551–618) and a GST-only control. All values are expressed as the number of released Zinc per molecule of protein, detected by change in PAR absorbance at 500 nm. One and two Zinc ions per molecule are shown as dashed lines.

To test for variability in our Alphafold predictions, ASXL2 PHD modelling was replicated several additional times (ten models in total) to determine whether a viable N-terminal Zinc-binding site could be predicted (Figures 3d, 3e). Zinc-binding was always modelled in the tetrahedral orientation at the ASXL2 PHD C-terminus (Figure 3e, Kuppuraj et al., 2009). In contrast, the position of chelating residues was varied at the ASXL2 PHD N-terminal Zinc-binding site (Figure 3d). All models placed Zinc in different positions (highlighted, opaque Zinc ions), in the square planar orientation, and some had Cys1400 contributing to Zinc binding (five chelating residues, 125% occupancy, Kuppuraj et al., 2009). Similarly, when AlphaFold models for the ASXL1, ASXL3 and *Drosophila* Asx PHD domains were generated, the N-terminal Zinc-binding site was always modelled poorly (Sup. Fig. 4, Kuppuraj et al., 2009). Given our difficulty in modelling the N-terminal Zinc binding site on the ASXL PHD, it appears that the domain may not bind to two Zinc ions (Aravind & Iyer, 2012), and is at least at the limits of the ability of Alphafold3 predictions.

An assay was subsequently performed to determine the number of Zinc bound by the ASXL PHD *in vitro*. Zinc ions were liberated from protein domains by denaturing samples in 8 M Guanidine-HCl prior to addition of 4-(2-pyridilazo) resorcinol (PAR) for Zinc detection (Doyle et al., 2019; Hunt et al., 1985). GST-fused versions of the human ASXL2 and *Drosophila* Asx PHD domains were tested and compared to the RING of Cellular Inhibitor of Apoptosis Protein 1 (cIAP1, positive control, two Zinc ions per molecule, Feltham et al., 2011) and GST alone (negative control, Sup. Fig. 5). The cIAP1 RING and GST control bound two and zero zinc ions on average, respectively. In contrast, the PHD domains of ASXL2 and Asx bound one Zinc ion per molecule (Figure 3f). Coupled with the AlphaFold models in Figures 3a-e and Sup. Fig. 3, this Zinc-binding data (Figure 3f) strongly suggests that the PHD domains of the human ASXL and *Drosophila* Asx proteins are not able to chelate Zinc at their N-terminus. Therefore, the ASXL PHD appears divergent from other binuclear treble-clef fingers (Dodd et al., 2004; Kaur & Subramanian, 2016; Li & Li, 2012; Matthews et al., 2009) and the domain may more accurately be described as a small, mono-nuclear Zinc knuckle. Given the lack of comparable functional domains, further biochemical studies were performed to determine the role of the ASXL PHD in PR-DUB complex function.

### The ASXL PHD forms a stable protein complex and composite Zinc-binding site with the MBD domains of MBD5 and MBD6

Recently, Tsuboyama and colleagues observed that the PHD domains of ASXL1–3 interact with MBD5 and MBD6, a set of previously-described PR-DUB interactors (Baymaz et al., 2014; Tsuboyama et al., 2022). MBD5 and MBD6 are the most poorly-characterised members of the Methyl-CpG-binding domain (MBD) family of proteins and have uniquely been shown not to bind methylated DNA (Laget et al., 2010). To explore their interaction with ASXL proteins further, we tested binding of the MBD domains of MBD5 (residues 18–81) and MBD6 (residues 17–81) to recombinant ASXL2 PHD domain protein (residues 1399–1435). Following expression in *E. coli*, MBD5 and MBD6 co-purified with GST-fused ASXL2 PHD (Figure 4a). We found that the ASXL2 PHD domain produced a higher yield with MBD6, than in the corresponding co-purification with MBD5 (Figure 4a). Given the MBD domains of MBD5 and MBD6 have high sequence similarity (68% identity), only the ASXL2 PHD-MBD6 protein complex was studied in subsequent experiments.

**Figure 4.**
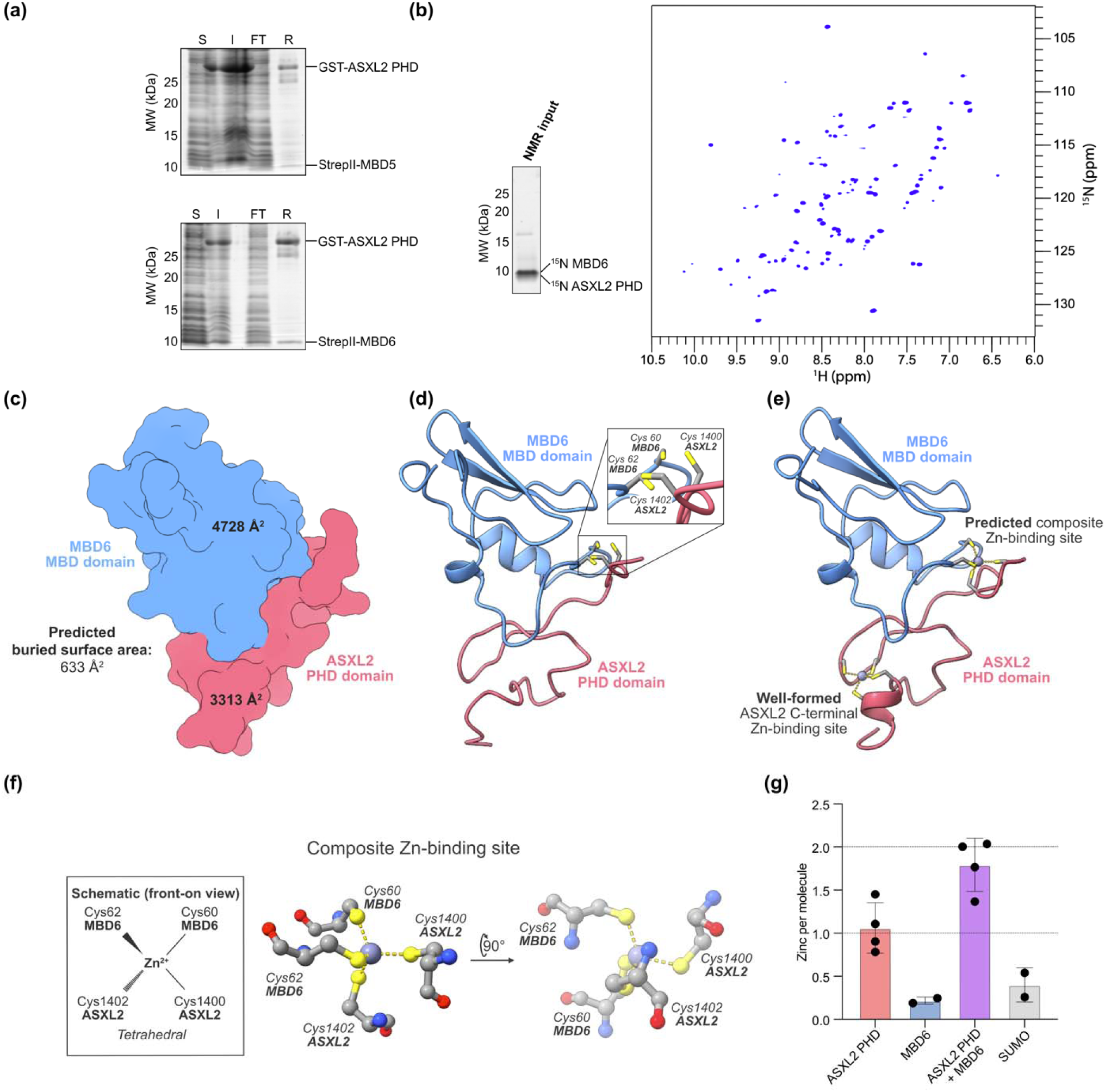
The ASXL PHD forms a composite Zinc-binding site with MBD5 and MBD6. **(a)** Co-purification of the MBD domains of MBD5 (residues 18–81) and MBD6 (residues 17–81) with the GST-fused PHD domain of ASXL2 (residues 1399–1435). Soluble (S), insoluble (I), and flow-through (FT) samples were collected and run alongside a single sample of resuspended GSH resin (R) on SDS-PAGE. A protein molecular weight marker was run alongside (MW, left) with sizes shown in kDa. **(b)** A co-expression construct was used to purify un-tagged ASXL2 PHD and MBD6 as a protein complex in ^15^NH_4_Cl minimal medium to yield uniformly ^15^N isotope-labeled protein. SDS-PAGE of the input protein for NMR is shown (left), alongside a solution ^1^H^15^N HSQC spectrum of the 42.5 μM complex sample. **(c)** Surface representation of the ASXL2 PHD (residues 1399–1435, pink) and MBD6 MBD domain (residues 17–81, blue) complex modeled in AlphaFold3. The total surface area of either protein domain and the buried surface area of the protein interface are labeled. **(d)** Cartoon representation of the ASXL2 PHD (residues 1399–1435, pink) and MBD6 MBD domain (residues 17–81, blue) complex modeled in AlphaFold3. Zinc-chelating residues that cluster near each other are shown in the enlarged region (inset, labeled). **(e)** Cartoon representation of the ASXL2 PHD (residues 1399–1435, pink) and MBD6 MBD domain (residues 17–81, blue) complex modeled in AlphaFold3 to bind two Zinc ions. The predicted, well-formed C-terminal and composite Zinc-binding sites are labeled. **(f)** The residues that contribute to the tetrahedral, composite Zinc-binding site that is predicted to form between the PHD and MBD domains of ASXL2 and MBD6, shown as in Figures 3b, 3c. **(g)** Zinc-binding assay comparing samples of the isolated ASXL2 PHD (residues 1399– 1435, N-terminal SUMO3 tag) and MBD6 MBD domains (residues 17–81, N-terminal Strep tag) with their co-purified complex (ASXL2 PHD + MBD6). A sample of the isolated SUMO3 fusion protein was included to control for variability of the Zinc assay.

A co-expression construct was used to generate ^15^N uniformly isotope-labelled ASXL2 PHD-MBD6 for characterisation by two-dimensional solution NMR spectroscopy (Figure 4b). The ^1^H^15^N HSQC spectra revealed the presence of about 95 backbone amide peaks, which is in agreement with the overall protein complex. The well-dispersed peaks and the lack of intense resonances in the unstructured central region suggest that both the ASXL PHD and MBD6 MBD domain were well-folded, and form a tight, stable protein complex *in vitro*. Consequently, we predicted the structure of the ASXL2 PHD-MBD6 protein complex using AlphaFold3 (Figure 4c, 4d, Sup. Fig. 6). The predicted protein-protein interface had a large buried surface area (633 Å^2^) relative to the total surface area of either domain (3313 Å^2^ for the ASXL2 PHD, 4728 Å^2^ for the MBD domain of MBD6, Figure 4c). Furthermore, we noticed residues Cys60 and Cys62 on MBD6 were positioned near Cys1400 and Cys1402 at the N-terminus of the ASXL2 PHD (Figure 4d). Given the proximity of Zinc-chelating residues within the modelled ASXL2 PHD-MBD6 complex, we considered the possibility that a shared, composite Zinc-binding site may exist at the interface between the two proteins (Figure 4d). Further AlphaFold modelling supported this observation, where two Zinc ions were modelled onto the complex in Figure 4e (Predicted Aligned Error (PAE) plot and pLDDT score colouring in Sup. Fig. 6). A tetrahedral Zinc-chelating site was predicted at the region described (labelled), with contributing residues from both interacting proteins (schematic, stick representation in Figure 4f).

Zinc-binding was subsequently compared between the isolated ASXL2 PHD-MBD6 complex and the constituent proteins, using the PAR-based Zinc assay established in Figure 3f. In contrast to the ASXL2 PHD (1.06 Zinc ions released per molecule, on average) and MBD6 MBD domain proteins by themselves (0.217 Zinc ions released per molecule, on average), the ASXL2 PHD-MBD6 complex released around two Zinc ions per molecule, on average (1.79, Figure 4f). Accordingly, the ASXL PHD likely assembles a composite Zinc-binding site when bound to the MBD domain of MBD6 (Figure 4e). Given the apparent flexibility of the ASXL N-terminus in earlier AlphaFold models (Figure 3), the composite Zinc-binding site likely increases the stability of the ASXL PHD N-terminus and mediates stable binding to MBD5 and MBD6.

### Misannotated domains mediate ASXL-MBD association

It is worth noting that the ASXL-MBD interaction is mediated by two domain types functioning non-canonically relative to their annotation. As outlined in this work, the ASXL-PHD domains do not appear to have capacity to bind histone marks, and instead function as protein-protein interaction domains. In parallel, the MBD domains of MBD5/6 are two members of the MBD family that are not capable of binding to methylated DNA. Mapping sequence differences on the PHD-MBD structural model generated in this work shows how these adaptions are mediated (Figure 5). Notably missing from the MBD domains of MBD5 and MBD6 are the amino acid residues of MBD1-4 loop L1 (labelled ‘L1 loop’, Figure 5). Upon binding to DNA, loop L1 undergoes major structural rearrangements, with seven of the nine L1 residues contributing to an interface with the DNA major groove (Ohki et al., 2001). MBD1-4 loop L1 residues have also been shown to specifically contact the Methyl-CpG site, involving residues absent from MBD5 and MBD6 (Ohki et al., 2001).

**Figure 5.**
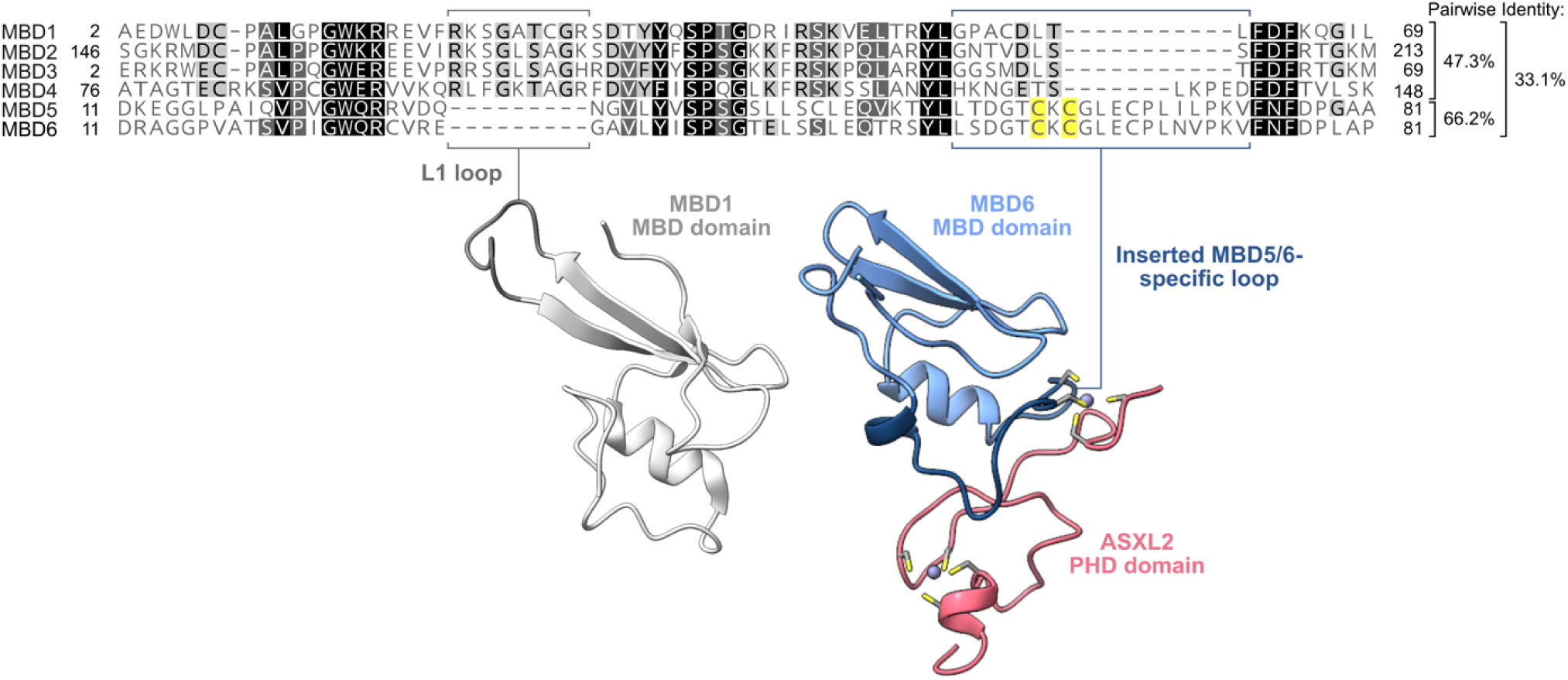
Multiple sequence alignment of MBD domains between members of the Human MBD protein family. The MBD domains of human MBD1–6 aligned by sequence similarity, and grouped by function (MBD1–4 all bind methylated DNA; MBD5–6 have non-canonical roles). A pairwise identity matrix (right) compares sequence similarity within functional groups, and between them. The structure of the MBD1 MBD domain (PDB: 6D1T) is shown, with the conserved methyl-DNA binding loop L1 highlighted in dark grey. Our AlphaFold3 model of the ASXL2 PHD-MBD6 protein complex is shown alongside (Figure 4), with the ASXL PHD interacting loop on MBD6 in dark blue. Conserved residues are highlighted on the multiple sequence alignment, with totally conserved residues in black, functionally conserved residues in grey, and the Zinc-binding Cysteine residues on MBD5/6 in yellow.

The MBD domains of MBD5 and MBD6 contain a specific insert between the helix α1 and the C-terminal hairpin loop that we and others show binds to the ASXL PHD (labelled ‘inserted MBD5/6-specific loop’, Figure 4, 5, Ohki et al., 2001; Tsuboyama et al., 2022). In conventional MBD domains, this loop is notably shorter (Figure 5, Ohki et al., 2001). In conjunction with this insertion, the cysteine residues Cys60 and Cys62 (MBD6 numbering) are conserved in only MBD5 and MBD6, but not conventional MBD family members (Figure 5). It is notable that the Zinc-chelating residues at the interface between the ASXL PHD, MBD5 and MBD6 all are conserved through to *Drosophila*, where the protein orthologs Asx (residues 1634–1669) and Six-banded (residues 238–308, homolog of MBD5 and MBD6 uncovered in Tsuboyama *et al*., 2022) are predicted to interact in the same orientation as in humans, shown in additional models we generated in AlphaFold3 (Sup. Fig. 8). Therefore, close co-operation between ASXL proteins and MBD5/6 equivalents appears to be an evolutionarily-conserved link in the epigenetic regulation of metazoans.

## Discussion

Truncation of the ASXL proteins is common in inherited and acquired disease, where loss of the ASXL C-terminus alters PR-DUB stability and deubiquitinase activity (Asada et al., 2018; Balasubramani et al., 2015; Inoue et al., 2016). In the current study, we examined biochemical functions of the ASXL PHD to determine how loss of the domain may contribute to the disease phenotype. Characterisation of the ASXL PHD was initially limited to identification of the Cys^4^-His-Cys^3^ Zinc-chelating motif that is typical to other PHD domains (Aravind & Iyer, 2012). This putative classification implied that the ASXL PHD may coordinate targeting of the PR-DUB to chromatin (Aravind & Iyer, 2012; Li & Li, 2012; Sanchez & Zhou, 2011; Slama & Geman, 2011). To the contrary, we have shown that the ASXL PHD domain is missing a complement of sequence features that are integral to histone methyl-binding (Figure 1). ITC (Figure 2a) and an ELISA (Figure 2b) of histone methyl modifications were subsequently used to demonstrate that the ASXL PHD is not able to bind common histone marks *in vitro*.

The ASXL PHD has previously been suggested to be atypical, where Aravind and Iyer identified that the domain lacks the bilobed brace that is required to bind the end of the histone H3 N-terminal tail. This feature was hypothesised to allow the ASXL PHD to bind histone modifications further up the histone tail, like H3K27me3 (Aravind & Iyer, 2012). Contrastingly, peptide pulldown assays performed by Park and colleagues showed broad specificity for the ASXL2 PHD to all of unmodified histone H3, H3K4me2 and H3K4me3 (Park et al., 2016). PHD domains typically are sensitive and selective to particular histone modification states, meaning the observed specificity to a broad range of modifications is unusual (Li & Li, 2012; Sanchez & Zhou, 2011; Slama & Geman, 2011). According to the current study, prior characterizations of the ASXL PHD overstate the ability of the domain to bind histone methyl marks. Instead, the ASXL PHD appears to have an atypical role in PR-DUB complex regulation. In support of an unusual function for the ASXL PHD, the fold of the domain is irregular (Figure 3). Most notably, the ASXL PHD is able only to chelate a single Zinc ion even though the domain sequence contains various potential residues that could chelate two Zincs (Figure 3). As a result, the N-terminus of the isolated ASXL PHD was variably-modelled in AlphaFold3 and appears flexible (Figure 3d).

During the course of this work, Tsuboyama *et al*. uncovered that the PHD domains of ASXL1–3 incorporate MBD5 and MBD6 into the PR-DUB complex (Tsuboyama et al., 2022). We could recapitulate binding of MBD5 and MBD6 with recombinant ASXL2 PHD (Figure 4a) and show that the protein complex is folded and stable in solution (Figure 4b). AlphaFold modelling was later used to predict the protein interface, where a large buried surface area relative to the total protein surface area was suggestive of a tight interaction (Figure 4c). Furthermore, Zinc-chelating residues on the ASXL PHD, MBD5 and MBD6 were shown to be proximal (Figure 4d, 4e, 4f) and to chelate an additional Zinc ion (Figure 4g) when in complex with one-another. Various other protein complexes bind Zinc at their interface to improve stability, and to provide sensitivity to intracellular concentrations of Zinc (Kocyła et al., 2021).

A mechanistic understanding of MBD5 and MBD6 protein function is elusive. It has been shown that MBD5 and MBD6 bind the ASXL PHD domain in a mutually-exclusive fashion (Baymaz et al., 2014; Tsuboyama et al., 2022) and that the MBD proteins co-localise with key components of the PR-DUB to chromatin (Tsuboyama et al., 2022). Furthermore, MBD5 and MBD6 each bind chromatin independently of the PR-DUB (Baymaz et al., 2014; Tsuboyama et al., 2022). MBD6, for instance, can independently co-localise to double-stranded DNA damage during the DNA damage response (Baymaz et al., 2014). Our current study further brings MBD5 and MBD6 into focus as key interactors of the ASXL PHD. We anticipate that future structural and biochemical studies will enable further insight into the ASXL PHD-MBD protein axis and clarify the functions of MBD5 and MBD6 within the PR-DUB complex. It may be interesting to investigate whether other protein interactors can bind the ASXL PHD in a fashion that is complementary to, or independent of, MBD5 and MBD6.

In summary, this study provides novel insight into the functions of the ASXL PHD domain. We show that the ASXL PHD is missing key functional residues and is unable to bind common histone modifications *in vitro*. Extensive AlphaFold modelling and validation through Zinc-binding assays demonstrate that the structure of the ASXL PHD is unusual but supported in complex with MBD5 and MBD6 through binding of an additional Zinc ion at the domain N-terminus. The latter finding highlights that MBD5 and MBD6 are key ASXL PHD interactors that likely contribute to ASXL truncation-driven disease.

## Acknowledgements

We thank all members of the Mace laboratory group, especially Abigail Burgess and Sam Jamieson for their helpful discussions and assistance during this project. We further acknowledge support for solution NMR measurements by Dr Vanessa Kay Morris. Funding support was provided by a project grant from the Health Research Council of New Zealand, a University of Otago Postgraduate Scholarship (to C.J.R.) and a University of Otago Master’s Research Scholarship (to A.R.W).

## Author contributions

C.J.R. performed experiments to characterise the isolated ASXL PHD domain. A.R.W. studied the interaction of the ASXL PHD with MBD5 and MBD6. C.G. performed and analysed the NMR experiment. P.D.M. obtained funding, supervised the study and wrote the manuscript together with C.J.R. All authors contributed to editing the manuscript.

