## Supplemental Data for "Unconventional structure and function of PHD domains from Additional Sex Combs-like proteins"

**Supplementary Figures**

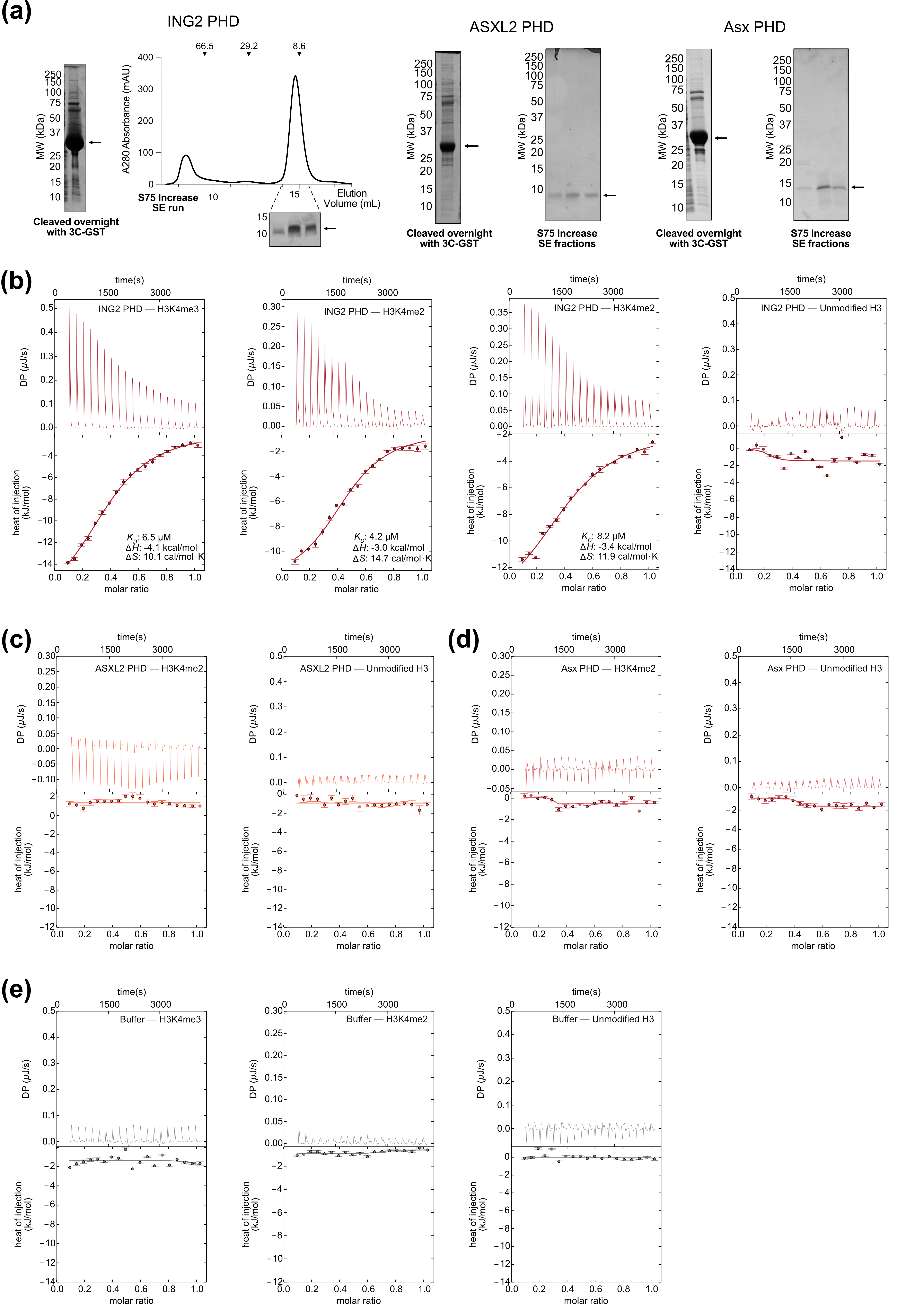

**Supplementary Figure 1.** Additional data for the Histone H3 peptide ITC analysis. (a) Example purifications of the ING2 (left, residues 199–261), ASXL2 (centre, residues 1375–1435) and *Drosophila* Asx (right, residues 1610–1668) PHD domains for ITC (arrows point to protein bands of interest).Protein domains were expressed fused to a GST tag, purified and cleaved overnight before being run on a S75 Increase 10/300 SE chromatography column into a buffer of 300 mM NaCl, 20 mM HEPES, 0.5 mM TCEP. Protein molecular weight markers (MW) are labelled on the Coomassie-stained SDS-PAGE gels; protein standards were run in the same buffer for SE chromatography and are labelled on the chromatogram (BSA, 66.5 kDa; Carbonic Anhydrase, 29.2 kDa; Ubiquitin, 8.6 kDa, arrowheads). (b) Heat change, measured by ITC, for interaction of the ING2 PHD domain (residues 199–261) with peptides of H3K4me3 (residues 1–10, replicate 2), H3K4me2 (residues 1–10, in duplicate) and unmodified histone H3 (residues 1–8). Key binding data are listed, where relevant. (c) Heat change, measured by ITC, for interaction of the ASXL2 PHD domain (residues 1375–1435) with peptides of H3K4me2 and unmodified histone H3. (d) Heat change, measured by ITC, for interaction of the *Drosophila* Asx PHD domain (residues 1610–1668) with peptides of H3K4me2 and unmodified histone H3. (e) Heat change, measured by ITC, for titration of histone H3 peptides against the buffer that all PHD domain proteins were prepared in (300 mM NaCl, 20 mM HEPES, 0.5 mM TCEP).

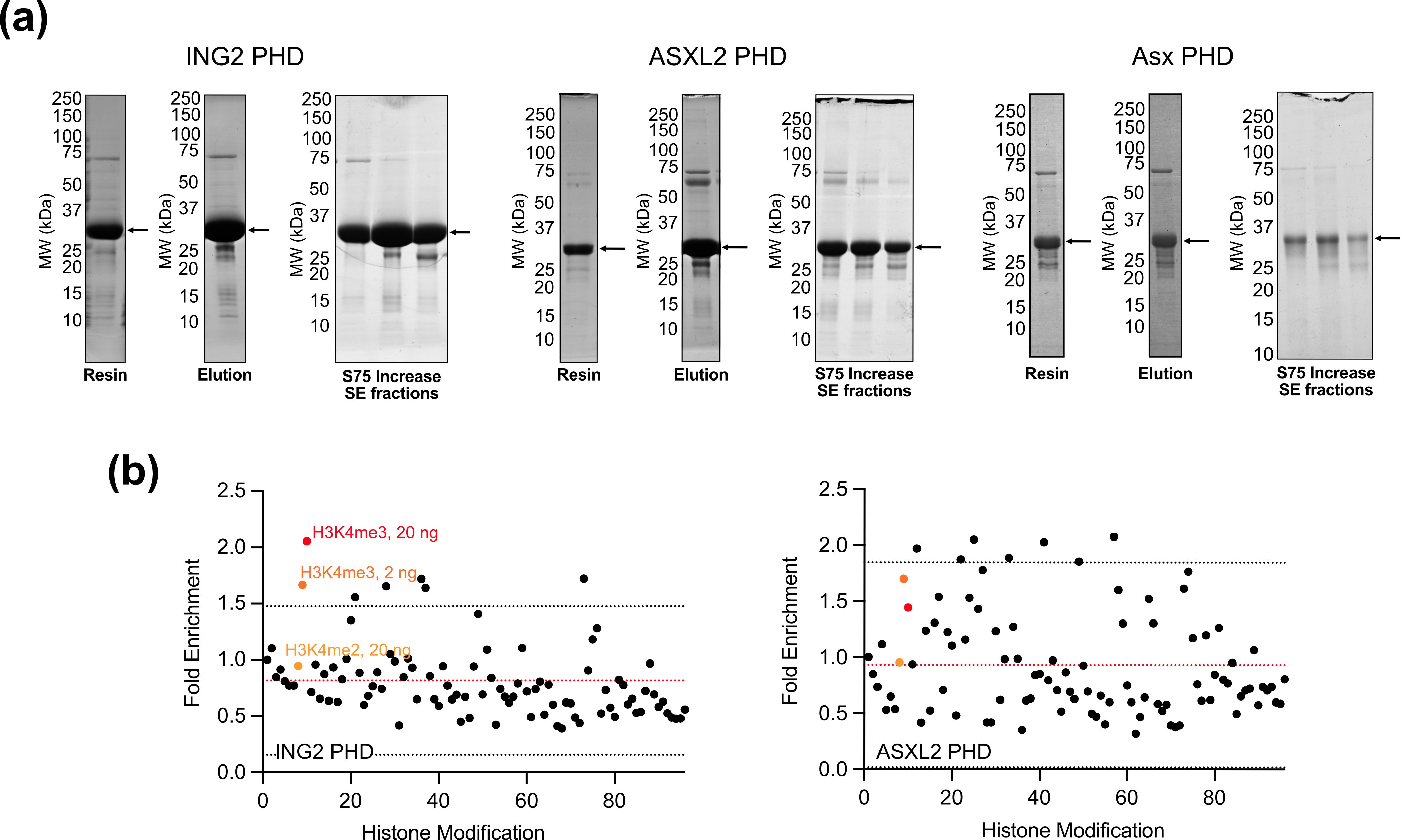

**Supplementary Figure 2.** Additional data for the histone H3 ELISA experiment. (a) Example purifications of the ING2 (left, residues 199–261), ASXL2 (centre, residues 1375–1435) and *Drosophila* Asx (right, residues 1610–1668) PHD domains for the histone H3 ELISA experiment (arrows point to protein bands of interest). Protein domains were expressed fused to a GST tag, purified, eluted from GSH resin and run on a S75 Increase 10/300 SE chromatography column into a buffer of 300 mM NaCl, 20 mM HEPES, 2 mM DTT. Protein molecular weight markers (MW) are labelled on the Coomassie-stained SDS-PAGE gels. (b) Fold enrichment of protein signal collected in replicates of histone H3 ELISA experiments, titrated with samples of the ING2 (residues 199–261) and ASXL2 (residues 1375–1435) PHD domains. Single points correspond to each of the 96 peptides spotted on the ELISA plate, in the order displayed in Appendix Table 1. Values are derived from the fold-enrichment of HRP signal relative to blank. The mean (red) and two standard deviations from the mean (black) are dashed lines. Peptides of H3K4me2 (20 ng) and H3K4me3 (2, 20 ng) are highlighted yellow, orange and red, respectively.

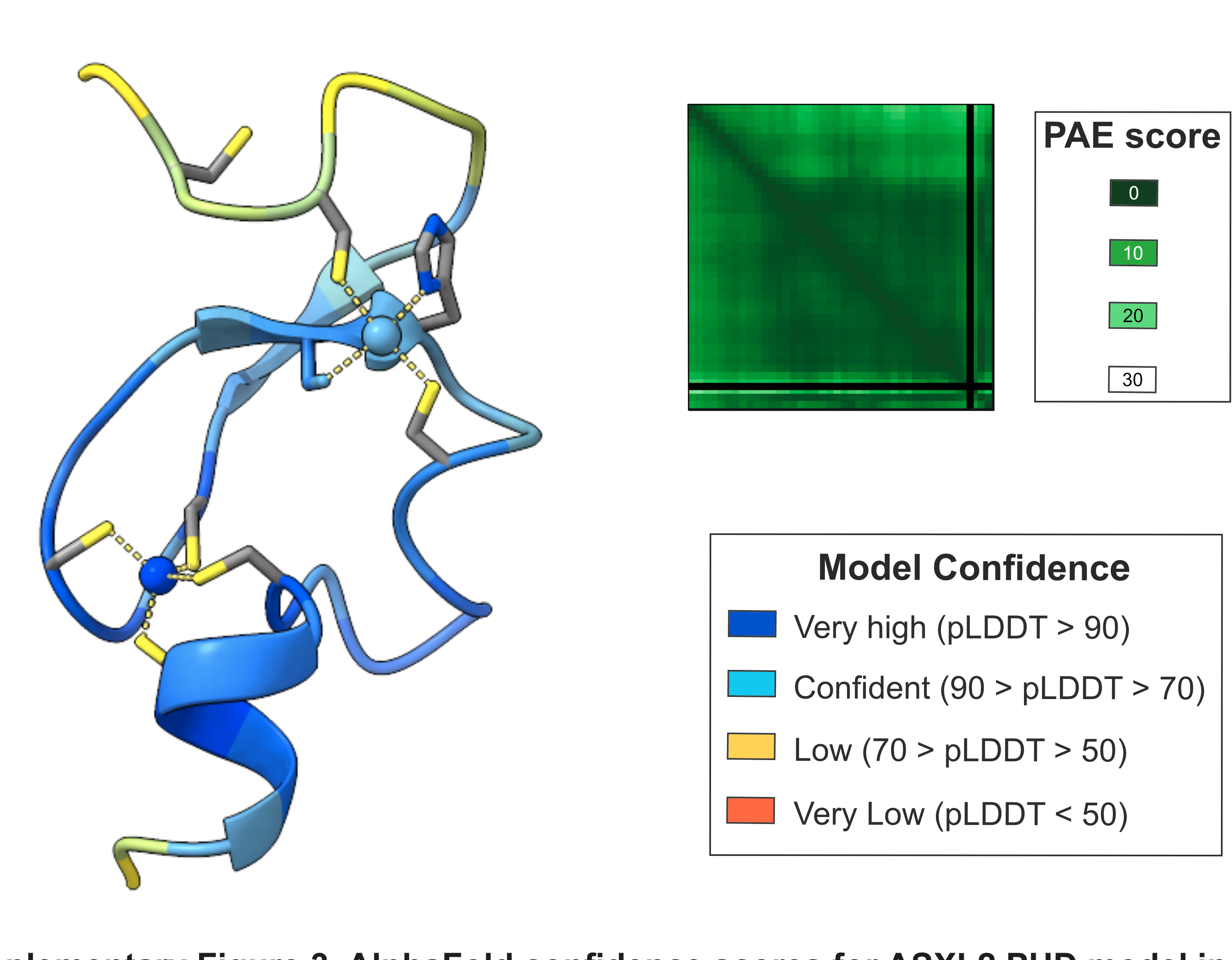

**Supplementary Figure 3.** AlphaFold confidence scores for ASXL2 PHD model in Figure 3a. AlphaFold3-derived structure of the ASXL2 PHD domain (residues 1399–1435) with two Zinc ions bound, coloured by predicted local distance difference test (pLDDT) score ('Model Confidence' key inset). The pLDDT score is a per-residue measure of local confidence. The Predicted Aligned Error (PAE) plot for the structure is shown alongside, coloured according to the PAE score key (inset).

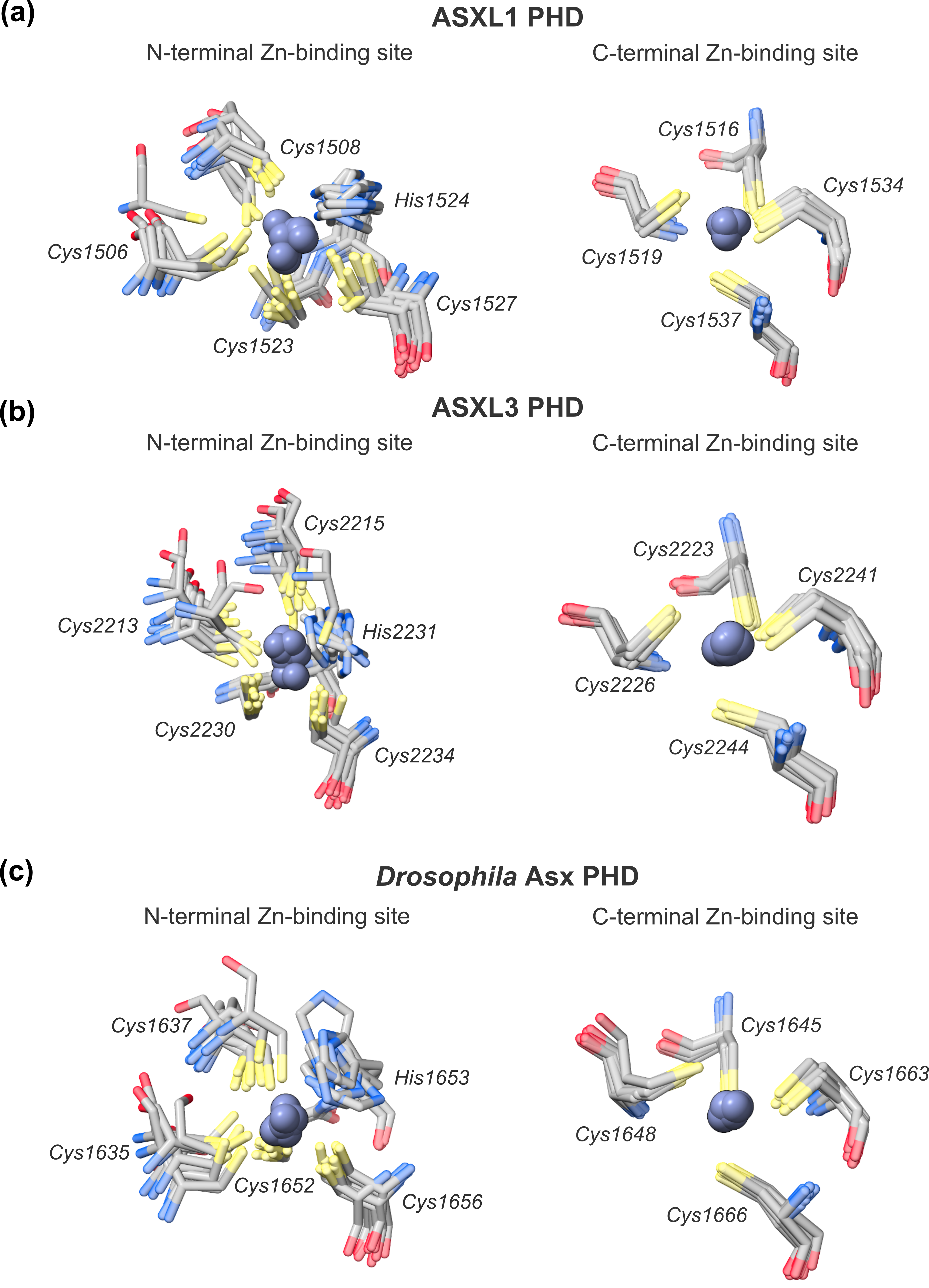

**Supplementary Figure 4.** Variable AlphaFold3 modelling of Zinc-binding sites between isoforms of ASXL1–3 and *Drosophila* Asx. (a) The N- and C-terminal Zinc-binding sites of the ASXL1 PHD, modelled ten times in AlphaFold3 (residues 1505–1541). Relevant, Zinc-chelating residues are shown as translucent sticks. The variably-modelled position of Zinc within either site is highlighted (silhouettes, opaque). (b) The N- and C-terminal Zinc-binding sites of the ASXL3 PHD, modelled ten times in AlphaFold 3 (residues 1399–1435), shown as in Sup. Fig. 4a. (c) The N- and C-terminal Zinc-binding sites of the *Drosophila* Asx PHD, modelled ten times in AlphaFold 3 (residues 1399–1435), shown as in Sup. Fig. 4a.

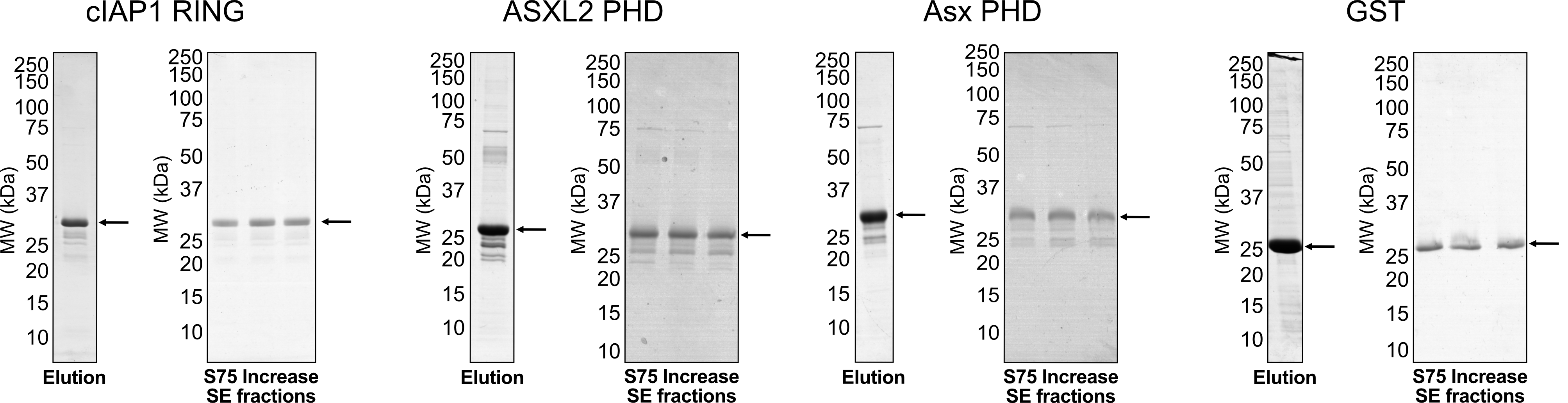

**Supplementary Figure 5.** Example purification of proteins for initial Zinc-binding experiments. Example purifications of the cIAP1 RING (left, residues 551–618), ASXL2 (centre left, residues 1375–1435) and *Drosophila* Asx (centre right, residues 1610–1668) PHD domains fused to GST, and GST alone (right), for the initial set of Zinc-binding experiments (arrows point to protein bands of interest). Protein domains were expressed fused to a GST tag, purified, eluted from GSH resin and run on a S75 Increase 10/300 SE chromatography column into a buffer of 300 mM NaCl, 20 mM HEPES. Protein molecular weight markers (MW) are labelled on the Coomassie-stained SDS-PAGE gels.

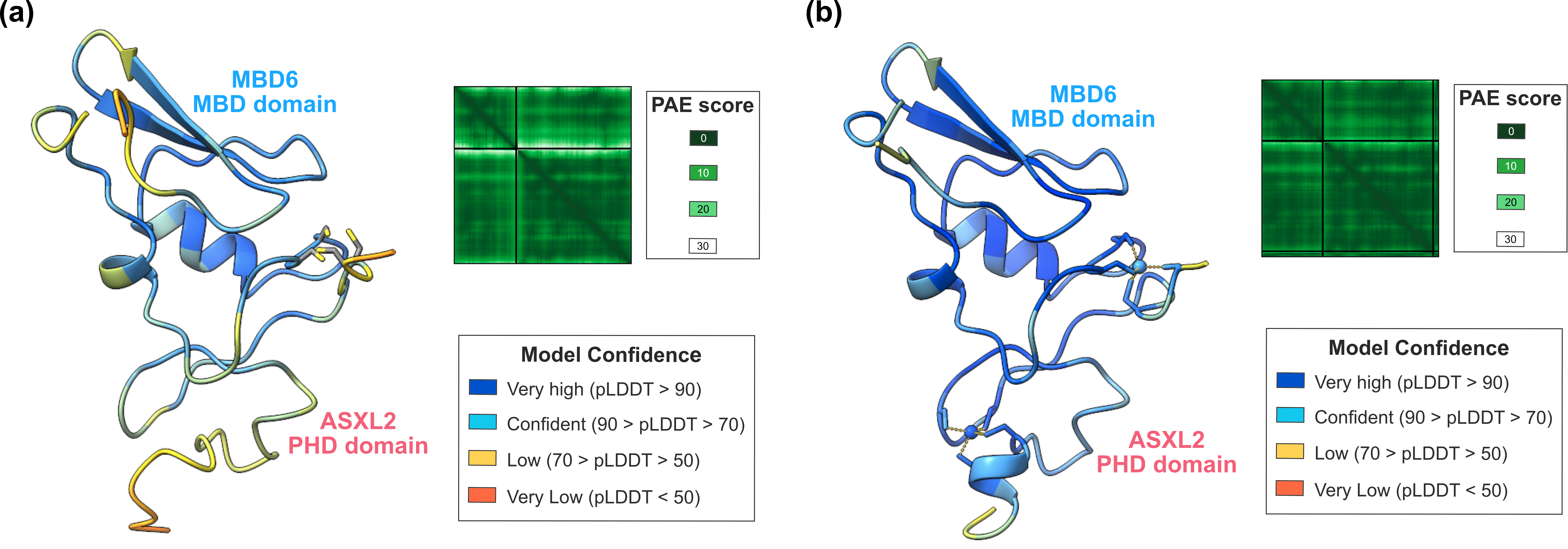

**Supplementary Figure 6.** AlphaFold confidence scores for ASXL2 PHD-MBD6 MBD domain complex models in Figures 4d, 4e. (a) Structure of the ASXL2 PHD (residues 1399–1435) - MBD6 MBD domain (residues 17–81) complex predicted using AlphaFold3 (Figure 4d), displayed in the same organisation as Supplementary Figure 3 (coloured by pLDDT score, PAE plot shown alongside). (b) Structure of the ASXL2 PHD (residues 1399–1435) - MBD6 MBD domain (residues 17–81) complex with two Zinc ions bound, predicted using AlphaFold3 (Figure 4e), shown in the same organisation as Supplementary Figure 3 (coloured by pLDDT score, PAE plot shown alongside). The overall confidence of the prediction increased with Zinc bound, relative to Sup. Fig. 6a.

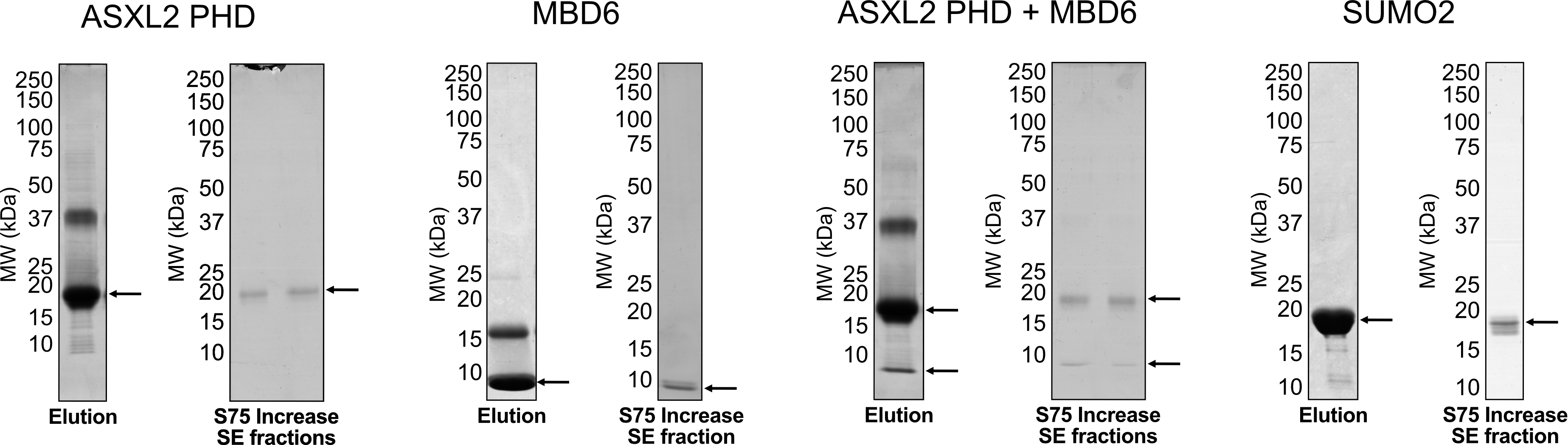

**Supplementary Figure 7.** Example purifications of the ASXL2 PHD (left, SUMO2-fusion, residues 1399–1435) and MBD6 MBD domains (centre left, StrepII-fusion, residues 17–81), ASXL2 PHD and MBD6 MBD domain co-expression vector (centre right, SUMO2- and StrepII-fusion, residues 17–81, 1399–1435), and SUMO3 alone (right), for the ASXL2 PHD-MBD6 complex Zinc-binding experiments (arrows point to protein bands of interest). The ASXL2 PHD, SUMO2 and co-expression proteins were purified using Nickel-affinity resin, eluted in a buffer containing imidazole. The StrepII-MBD6 protein was purified using Strep-Tactin®XT 4Flow® high capacity resin, eluted in 1x Buffer BXT. All proteins were purified on a S75 Increase 10/300 SE chromatography column into a buffer of 300 mM NaCl,20 mM HEPES. Protein molecular weight markers (MW) are labelled on the Coomassie-stained SDS-PAGE gels.

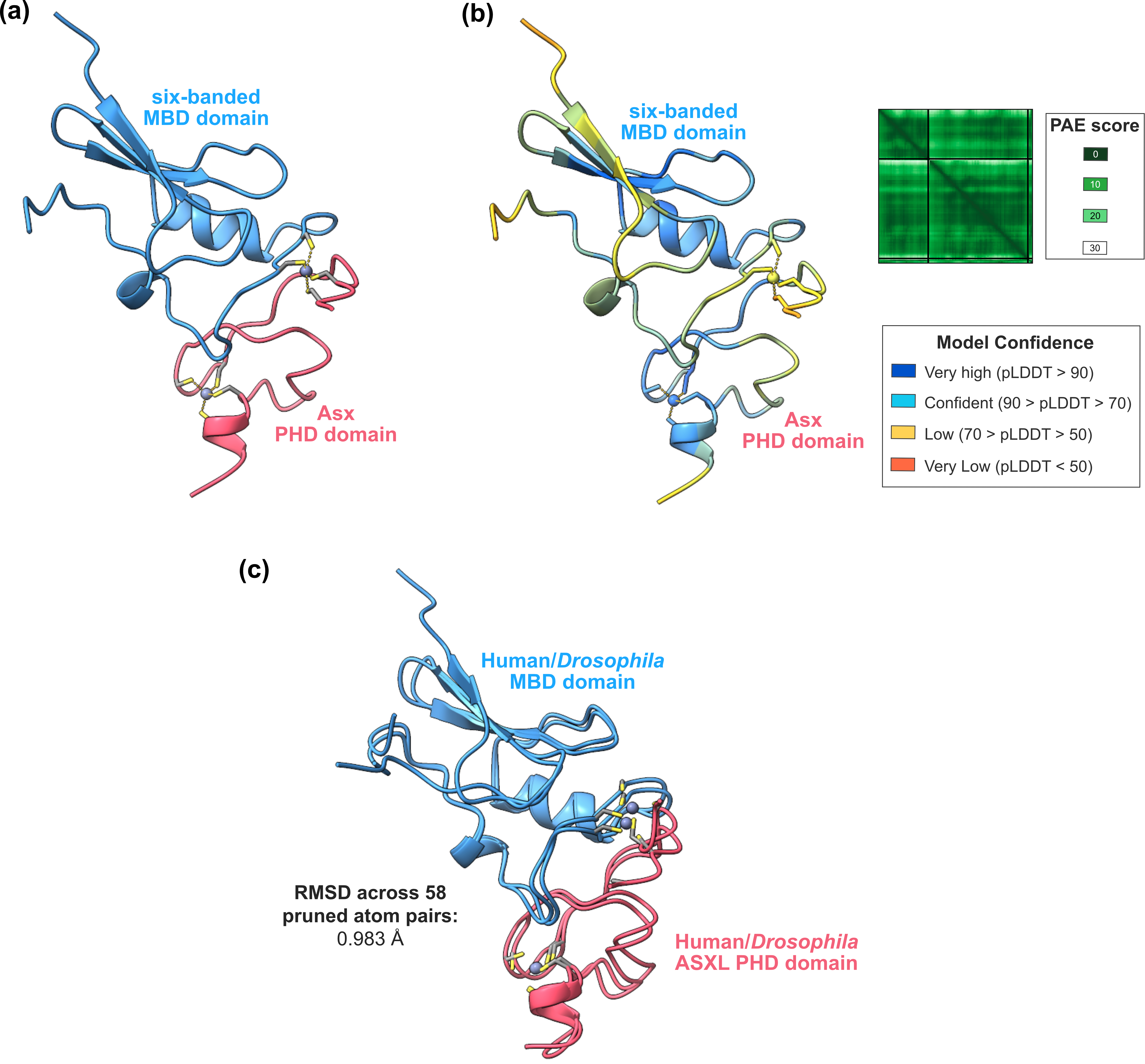

**Supplementary Figure 8.** Structure of *Drosophila* Asx and Six-banded in complex, predicted using AlphaFold3. (a) Structure of the Asx PHD (pink, residues 1634–1669) and Six-banded MBD domain (residues 238–308) complex with two Zinc ions bound, predicted using AlphaFold3. The proteins are shown in the same representation and colouring as for their human homologs in Figures 4d, 4e. (b) Supplementary Figure 6a coloured by pLDDT score (key inset) and with the PAE plot shown (key inset). (c) Alignment of the predicted human ASXL2 PHD-MBD6 complex with the *Drosophila* orthologs (Asx and Six-banded). The Root Mean Square Deviation (RMSD) of a pruned subset of atoms (58 of 66 total atom pairs) is shown. The low RMSD and highly-similar structural alignment suggest that the organisation of the Human and *Drosophila* protein complexes are similar.
